# Calcification and trophic responses of Mesophotic reefs to carbonate chemistry variability

**DOI:** 10.1101/2023.08.09.552600

**Authors:** Timothy J. Noyes, Rebecca Garley, Nicholas R. Bates

**Author notes:** **Correspondence:** Timothy J. Noyes.

## Abstract

Mesophotic coral ecosystems (MCEs) are extensions of adjacent shallow water coral reefs. Accessibility to these ecosystems is challenging due to their depth limits (∼ 30 – 150 m) and as a result, scientific knowledge of these reef systems is limited. It has been posited that the depth limits of MCEs diminish anthropogenic effects experienced by shallow reef systems. A lack of empirical measurements to date has made this hypothesis impossible to determine for mesophotic reef metabolism. The alkalinity anomaly technique was utilized to determine rates of net ecosystem calcification (NEC) and net ecosystem production (NEP) from 30, 40 and 60 m mesophotic reefs during a 15-month period. Seawater chemistry was determined to be chemically conducive for calcification (average aragonite saturation Ω_aragonite_ of 3.58, average calcite saturation Ω_calcite_ of 5.44) with estimates of NEC indicating these reef systems were net accretive and within global average values for shallow coral reefs (< 30 m). The strongest periods of calcification occurred in late summer and were coupled with strong autotrophic signals. These episodes were followed by suppressed calcification and autotrophy and in the case of the 60 m reefs, a switch to heterotrophy. Whilst there was variability between the three reefs depths, the overall status of the mesophotic system was net autotrophic. This determination was the opposite of trophic status estimates previously described for adjacent shallow reefs. Whilst there were periods of net dissolution, the mesophotic reef system was net accretive (i.e., gross calcification > gross CaCO_3_ dissolution). The measured inorganic carbon chemistry and estimates of NEC and NEP represent the first such biogeochemical measurements for MCEs. The values established by this study demonstrate just how close these understudied ecosystems are in terms of the known boundary thresholds for low saturation state reefs. Making predictions on how these ecosystems will respond to future climatic conditions, will require greater sampling effort over long times scales to decouple the environmental controls exerted on such ecosystems.

## 1 Introduction

Coral reefs are highly diverse marine ecosystems forming complex three-dimensional frameworks, generated through biogenically precipitated calcium carbonate (CaCO_3_) primarily produced by hermatypic corals, calcareous algae and foraminifera. Such calcification is responsible for ∼50% of net annual CaCO_3_ oceanic precipitation (Dubinsky and Stambler, 2011). Globally, coral reef ecosystems provide numerous socio-economic and ecological functions (Moberg and Folke, 1999; Sarkis et al., 2013). Their global decline through disturbance events, climate change and a myriad of anthropogenic activities have been well documented in the literature (e.g., (Wilkinson, 1999, 2008; Jackson et al., 2001, 2014; Pandolfi et al., 2003; Hughes et al., 2007; Hoegh-Guldberg, 2011; Tanzil et al., 2013). These declines have led to increased scientific interest in deeper reef ecosystems (e.g., mesophotic, generally deeper than 30 m) that may exhibit reduced susceptibilities and greater resistance to such environmental change (Glynn, 1996; Puglise et al., 2008; Bongaerts et al., 2010, 2017; Baker et al., 2016; Cinner et al., 2016; Loya et al., 2016; Semmler et al., 2017; Turner et al., 2017).

Deeper reef systems (generally deeper than 30 m with depths up to ∼ 150 m) termed “mesophotic coral ecosystems” (MCEs; Hinderstein et al., 2010) have a near-ubiquitous presence below shallow coral reefs (Bridge et al., 2013). The consensus within the research community is that MCEs are often extensions of shallow coral reef systems (Hinderstein et al., 2010; Pyle et al., 2019 and references therein) with common species shared between the two communities. Subdivisions of the MCE habitat based on biodiversity changes across depth have been proposed (Pyle et al., 2019 references therein). It is this commonality of species combined with the suggestion that MCEs could be buffered from anthropogenic influence (e.g., ocean warming and acidification) and natural disturbances (Bongaerts et al., 2013; Loya et al., 2016) that makes these ecosystems of particular scientific interest. Determining the levels of oceanographic connectivity between mesophotic and adjacent shallow-water reefs has become a key research priority across disciplines (e.g., biogeochemistry, marine spatial planning). Given that biological carbonates are the largest carbon reservoirs in the biosphere (i.e., aragonite, calcite, and magnesian calcite; Cohen, 2003), to truly comprehend the level of resilience coral reef systems have to anthropogenic and natural stressors, understanding the interactions and feedbacks that drive these biogeochemical processes (e.g., calcification rates, thermal regimes, flow dynamics) are fundamental requirements. Oceanic uptake of anthropogenic CO_2_ has led to significant changes in seawater chemistry (Orr et al., 2005) which in turn has raised concerns about the consequences to marine calcifiers (e.g., hermatypic corals, calcareous algae and foraminifera; Kleypas et al., 1999; Hoegh-Guldberg et al., 2007) to generate and maintain CaCO_3_ structures through a reduction in seawater pH and aragonite saturation state (Ω_aragonite_; Cyronak et al., 2018; Eyre et al., 2018). Of equal importance is the potential for rates of bioerosion and dissolution of CaCO_3_ structures (Andersson et al., 2009; Tribollet et al., 2009; Dove et al., 2013) to increase under the same conditions. The persistence of coral reefs is dependent on their ability to calcify and produce CaCO_3_ and maintain net positive accretion. At reef scale, net accretion of CaCO_3_ and external sediment supply (e.g., broken down framework, shells) must be greater than any loss by dissolution, transport, or erosion (Kleypas et al., 2001; Andersson and Gledhill, 2013; Eyre et al., 2018).

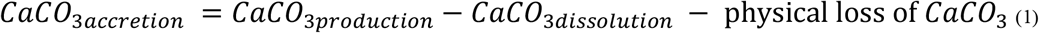

Both models and mesocosm studies on individual and community calcifiers have measured decreased rates in calcification (net accretion) and increases in CaCO_3_ dissolution ultimately transitioning to a state of net dissolution under projected atmospheric and seawater CO_2_ conditions (Pandolfi et al., 2011; Andersson and Gledhill, 2013; Lantz et al., 2014). However, there is considerable uncertainty as to when this inferred threshold may be crossed (Andersson and Gledhill, 2013). Rates of net coral CaCO_3_ production can be determined through chemistry-based methods (i.e., net ecosystem calcification [NEC] = gross calcification – gross CaCO_3_ dissolution) and can provide a top-down integrated measurement of the entire reef NEC (Courtney et al., 2016). Modification of CO_2_-carbonate chemistry of Bermudan mesophotic reefs will reflect the main biogeochemical processes occurring on the reef system (Bates, 2002, 2017; Bates et al., 2010; Andersson et al., 2014; Yeakel et al., 2015; Courtney et al., 2016). It is accepted that inorganic CaCO_3_ accretion occurs at seawater saturation state (Ω) >1 whereas the dissolution of CaCO_3_ occurs when Ω <1. However, biogenic reef carbonate dissolution has been determined to occur well above this expected thermodynamic transition value (Langdon et al., 2000; Bates et al., 2010; Andersson et al., 2014). Differences in open ocean source water CO_2_-carbonate chemistry are compared to water collected from mesophotic reefs thus allowing the determination of NEC (accretion or dissolution) and NEP (autotrophy or heterotrophy) consequently resolving if Bermudan MCEs are dominantly in a state of net calcification and whether the trophic status is in balance.Methodology

## 2 Methodology

### 2.1 Study Locations

Bermuda is located approximately 1000 km east south-east of the United States of America (Jones et al., 2012) in the northwest of the Sargasso Sea (Coates et al., 2013) and considered the northernmost reefs of the Atlantic (Spalding et al., 2001; Logan and Murdoch, 2011). Bermuda’s coral reef system transitions through a series of shallow patch reefs, shallow rim reefs, and terraced reefs (Logan, 1988; Logan and Murdoch, 2011) dropping quickly to deeper mesophotic reefs that surround the main reef platform (Goodbody-Gringley et al., 2019). Using the 30 – 150m contours in a Bermuda 1 arc-second sea level digital elevation model (Sutherland et al., 2014) as a proxy for the extent of the mesophotic zone surrounding Bermuda, the system covers 76 km^2^ which equates to 8% of habitats < 30 m (908 km^2^). All physiographic reef zones are influenced by the offshore waters from the Sargasso Sea (Steinberg et al., 2001; Bates, 2017). The study was performed on three mesophotic reef locations between August 2017 and October 2018 (Figure 1). The choice of locations were defined a posteriori following the findings of a Darwin Plus (DPLUS001) lionfish control initiative project completed in 2015. At each location, a grid of six sites approximately ∼ 350 m apart were arranged in a pattern of three shallow (30 ∼ 40 m) and three deep sites (60 m). The benthic community composition followed the pattern reported by Fricke and Meischner (1985) and expanded upon in the review of Bermuda’s mesophotic reefs by Goodbody-Gringley et al. (2019). Moving in a seaward direction, shallow mesophotic sites (∼30 m depth) were dominated by hermatypic scleractinian corals whilst the deeper sites exhibited greater benthic heterogeneity as macroalgae (Stefanoudis et al., 2019) and rhodolith beds became more dominant.

**Figure 1.**
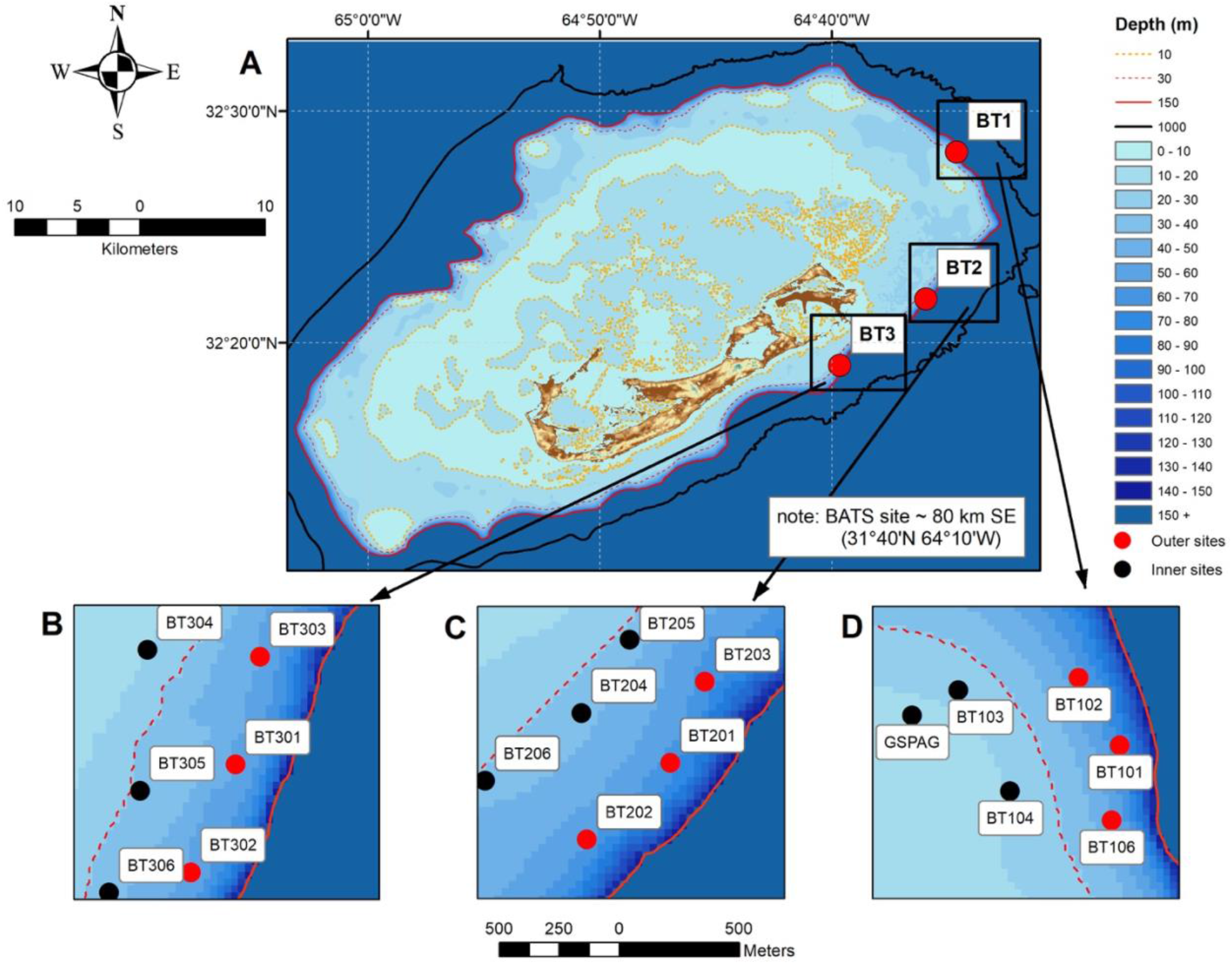
**(A)** Bathymetric map of Bermuda illustrating the main study locations (red circles) and proximity to land and open ocean, 10 m contour line in orange, 30 m contour dashed red line, 150 m contour solid red line, 1000 m contour solid black line. **(B)** location BT3 showing spatial orientation of ∼ 30 - 40 m (black circles), 60 m (red circles). **(C)** location BT2 showing spatial orientation of ∼ 30 - 40 m (black circles), 60 m (red circles). **(D)** location BT1 showing spatial orientation of ∼ 30 - 40 m (black circles), 60 m (red circles). Note, the red dashed line represents the ∼ 45 m contour for this location only.

### 2.2 Seawater Carbonate Chemistry Determination

Carbon chemistry samples were collected on an ad hoc basis between August 2017 and October 2018 using a 12-liter water sampler bottle (Standard Model 110B, OceanTest Equipment, Inc.) at ∼2 m above the benthos (Bates et al., 1996; Dickson et al., 2007). Comparative offshore samples were collected monthly as part of the Bermuda Atlantic Time-series Study (BATS; Bates et al., 2012). Samples for dissolved inorganic carbon (DIC) and total alkalinity (TA) were drawn into clean 300-ml Kimax glass sample bottles and fixed with 100 μl of saturated mercuric chloride (HgCl_2_) solution to prevent biological alteration. Dissolved inorganic carbon was analysed using an Automated InfraRed Inorganic Carbon Analyzer (AIRICA, Marianda Inc) or by coulometric technique on a VINDTA system (Versatile INstrument for the Determination of Total inorganic carbon and titration Alkalinity, VINDTA 3C, Marianda Inc). DIC is defined as (Dickson, Sabine and Christian, 2007):

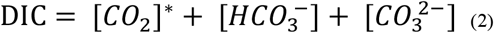

All reef TA samples were analysed via open-cell potentiometric titration with HCl of approximate normality of ∼0.1 and ionic strength of ∼0.7 using the VINDTA 3S system (Marianda Inc). The offshore BATS TA samples were analysed on a VINDTA 2S system (Marianda Inc) with similar solutions. The VINDTA2S system is the previous model of VINDTA 3S system and although it performs the same analytical function, BATS samples are preferentially run on this machine to maintain continuity, but with no demonstrable difference between analytical systems. TA is typically defined as:

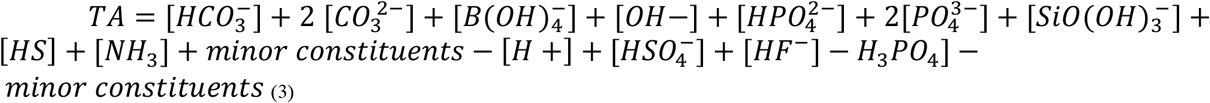

Seawater certified reference materials (CRMs; prepared by A.G. Dickson, Scripps Institution of Oceanography; http://www.dickson.ucsd.edu) were used to ensure the precision and accuracy of both DIC and TA values (typically ±1 to 2 μmoles kg^-1^). For both reef and BATS measurements, seawater pH, pCO_2_ and Ω were calculated using CO2SYS (Lewis and Wallace, 1998) from measured DIC and TA at in situ salinity and temperature conditions using the K1 and K2 dissociation constants from Mehrbach et al. (1973) refitted by Dickson and Millero (1987).

### 2.3 Physical and biogeochemical parameters

Salinity samples were paired with all reef and BATS DIC and TA samples and collected in accordance with best practices (Knap et al., 1997). Samples were drawn into 250 ml clear borosilicate glass bottles with plastic screw caps. A plastic insert was used to form an airtight seal and stop sample evaporation. All samples were analysed on a Guildline AutoSal 8400A laboratory salinometer (± 0.002) following the manufacturers recommendations for standard practices. Salinity measurements were calibrated against IAPSO standard seawater (Ocean Scientific, UK) to give a precision of ± 0.001-0.002 Practical Salinity Units (PSU). In situ temperatures were measured with an ONSET HOBO Water Temp Pro v2 (accuracy ±0.2°C between -40°C and 70°C). Inorganic nutrient samples were collected on an ad hoc basis (Table 1) at the central shallow and deep sites per each location only (Figure 1). Samples were collected following best practices (Knap et al., 1997), filtered through a 0.8 μm NucleporeTM filter (Whatman®) into prewashed 60 ml amber bottles (Nalgene® HDPE) stored on ice and immediately frozen on return to BIOS, prior to shipping to the Woods Hole Oceanographic Institution Nutrient Analytical Facility. All samples were analysed on a SEAL Analytical AA3 HR Auto Analyzer using U.S. Environmental Protection Agency methods for ammonium (method G-171-96, detection limit 0.015 μmoles L^-1^), nitrate + nitrate (method G-172-96, detection limit 0.040 μmoles L^-1^), silicate (method G-177-96, detection limit 0.030 μmoles L^-1^), and phosphate (G-297-03, detection limit 0.009 μmoles L^-1^).

**Table 1.** Summary of the number of physico-chemical parameters measured at three reef zones.

| Reef zone | Temperature | Salinity | DIC | TA | NH<sub>4</sub><sup>+</sup> | NO<sub>2</sub><sup>-</sup> ± NO<sub>3</sub><sup>-</sup> |
| --- | --- | --- | --- | --- | --- | --- |
| 30 m | 38 | 38 | 38 | 38 | 17 | 17 |
| 40 m | 41 | 41 | 41 | 41 | 17 | 17 |
| 60 m | 98 | 98 | 98 | 98 | 33 | 33 |
| Total | 177 | 177 | 177 | 177 | 67 | 67 |

| Reef zone | PO<sub>4</sub><sup>3-</sup> | Silicate |
| --- | --- | --- |
| 30 m | 17 | 17 |
| 40 m | 17 | 17 |
| 60 m | 33 | 33 |
| Total | 67 | 67 |

### 2.4 Determination of Net Ecosystem Calcification (NEC) and Net Ecosystem Production (NEP)

The determination of net ecosystem calcification (NEC) is based on the widely accepted alkalinity anomaly-water residence time technique (Smith and Key, 1975; Bates et al., 2010; Langdon et al., 2010; Andersson and Gledhill, 2013; Courtney et al., 2016; Bates, 2017). The relative changes in DIC and TA reflect biogeochemical partitioning of carbon between the inorganic and organic cycles (Suzuki and Kawahata, 2003). The changes in DIC and TA as a result of NEC change in a ratio 1:2 DIC:TA. Net calcification (NEC > 0) will reduce both DIC and TA which causes a lowering of pH and Ω_aragonite_.

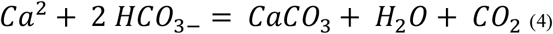

Offshore BATS samples are assumed to be representative of waters flowing onto Bermudan mesophotic reefs. The sampling regimes between the MCE sites and BATS were typically conducted within 1-2 weeks of each other (Bates, 2017). To minimise the influence of isopycnal lifting as water transitions onto Bermuda MCEs, comparative offshore data were selected based on salinity and temperature. To account for local evaporation and precipitation changes, all (i.e., MCE and BATS) TA and DIC were salinity normalised, nTA and nDIC respectively, to a mean measured salinity of MCE reefs of 36.67 (Courtney et al., 2021).

The method assumes any differences in total alkalinity (TA) between offshore and mesophotic reef seawater (i.e., nTA_offshore_ – nTA_MCE_) are a relative expression of MCE calcification and calculated as per the method of Langdon et al. (2010):

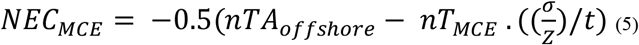

Where σ is the density of seawater, Z is the depth of water and t is the water residence time for the mesophotic reef. Water depths for MCE sites were measured using a vessel mounted depth sounder (Garmin GPSmap 441s/Aimar P79 50/200 kHz transducer). Sample depths for offshore (BATS) samples were recorded by a Sea-Bird SBE 911 CTD instrument package (SBE 9 underwater unit, SBE 11 Deck unit). Sites were categorized as 30 m, 40 m and 60 m as determined by average depth measurements recorded over the duration of the study. Seawater residence times for mesophotic sites were deemed to be 0.5 days based on hydrological modelling (R. Johnson unpublished data). All discrete reef level samples were collected from ∼ 2 m above the benthos, as such, any calcification and or dissolution signals (i.e., CaCO_3_ precipitation/dissolution) are estimated to be detected within a 5 m^3^ volume above the benthos. Rates of NEC_MCE_ are calculated in units of mmoles CaCO_3_ m^-2^ d^-1^ (or expressed as g CaCO_3_ m^-2^ d^-1^ using the molecular weight 100.09).

Net calcification (NEC > 0) draws down both DIC and TA causing a reduction in seawater pH and Ωaragonite. NEP alters DIC content of the water however, neither photosynthesis or respiration alter TA, therefore NEP_MCE_ is calculated as the difference between offshore BATS and onshore MCE samples as follows:

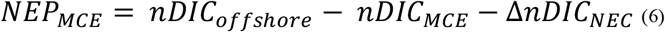

Air–sea CO_2_ gas exchange were deemed minor relative to the calculations of NEC and NEP (Bates, 2017). Rates of NEP_MCE_ are calculated in units of mmoles C m^-2^ d^-1^ (or expressed as g C m^-2^ d^-1^ using a molecular weight of 12).

### 2.5 Propagation of Uncertainty

Uncertainty for bottle NEC and NEP calculations were estimated using procedures outlines by Ku (1966). The uncertainty of measured DIC and TA (±1 μmoles kg^-1^) was obtained from routine measurement of CRMs (prepared by A.G. Dickson, Scripps Institution of Oceanography; http://www.dickson.ucsd.edu) with the DIC and TA samples.

## 3 Results

### 3.1 Physical and biogeochemical variability

In total 177 paired comparisons were made between mesophotic reefs and reference samples taken at the BATS site which when delineated to depth categories, equated to 30 m = 38, 40 m = 41, 60 m = 98 samples respectively (Table 1). Monthly benthic seawater temperatures exhibited seasonal variability across all study sites ranging from 19.6 – 27.5 ± 2.2 °C between winter and summer (Figure 2 A-C). The summer monthly climatology of 60 m reefs tended to be ∼ 3.5 to 4°C and ∼ 2.5 to 3°C cooler than the 30 m and 40 m reefs respectively. Short term deployments (∼ 12 days) of temperature loggers at 60 m sites recorded ∼4 °C daily variability. High degrees of thermal oscillation at mesophotic depths are known to occur (Wolanski et al., 2004; Colin, 2009; Colin and Lindfield, 2019) and have been attributed to internal waves. In the Pacific region of Micronesia, extreme daily thermal isolations at a 90 m depth (∼ 20 °C) were linked to a possible coupling of internal waves with a Rossby wave causing a deepening of the thermocline (Colin and Lindfield, 2019). Salinity across all mesophotic sites had a similar seasonal range to offshore values recorded at BATS 36.67 ± 0.09 g kg^-1^, however, samples from 2017 were slightly fresher than 2018 but still within the limits reported for the Bermuda environment (36.3 – 36.7; Coates et al., 2013; Yeakel et al., 2015). Inorganic nutrients concentrations were typically below 0.1 μmoles L^-1^ (oligotrophic water column; Table 2) and consistent with values previously published for Bermuda mesophotic reefs (Goodbody-Gringley et al., 2015). Approximately 50% of all inorganic nutrient measurements were below detection limits suggesting rapid uptake of dissolved inorganic nutrients and/or limited nutrient availability. Nutrient measurements were collected at a subset of sites (n = 67) over the duration of the study with a focus on the central 60 m site per location.

**Figure 2.**
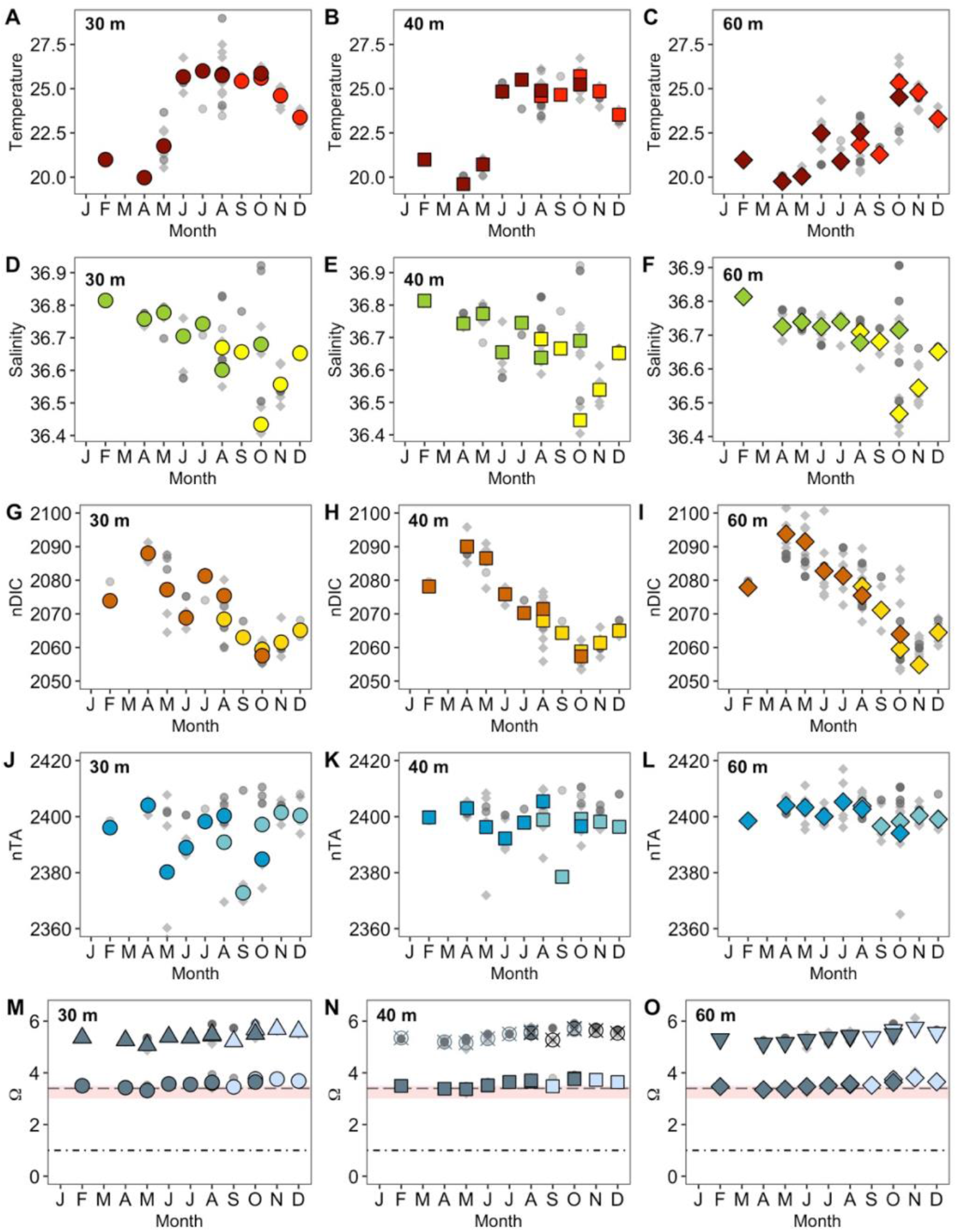
Mesophotic carbonate chemistry data at 30 m, 40 m and 60 m depths. **(A– O)**, BATS data (dark grey circles), MCE 2017 data (light grey circles), 2018 (grey diamonds). **(A - C)** Temperature °C (2017 red symbols, 2018 dark red symbols); **(D - F)** salinity PSU (2017 yellow symbols, 2018 green symbols); **(G - I)** nDIC (μmoles kg^-1^, 2017 orange symbols, 2018 dark orange symbols); **(J - L)** nTA (μmoles kg^-1^, 2017 blue symbols, 2018 dark blue symbols; **(M)** Ω_aragonite_ (2017 light blue circles, 2018 grey circles), Ω_calcite_ (2017 light blue triangles, 2018 grey triangles); **(N)** Ω_aragonite_ (2017 light blue squares, 2018 grey squares), Ω_calcite_ (2017 black hatched circles, 2018 light grey hatched circles); **(O)** Ω_aragonite_ (2017 light blue diamonds, 2018 grey diamonds), Ω_calcite_ (2017 light blue inverted triangles, 2018 grey inverted triangles). DIC, TA, Ω_aragonite_ and Ω_calcite_ have been salinity normalized to values of 36.67 g kg-1. The black dot - dash line depicts Ω_aragonite_ = 1, thermodynamically, dissolution is anticipated if Ω < 1; grey dashed line depicts Ω_aragonite_ = 3.4, transition from coral reef to non-reef coral community; pink shaded area Ω_aragonite_ 3.0 – 3.5 defined as the global limit for reef development.

**Table 2.**
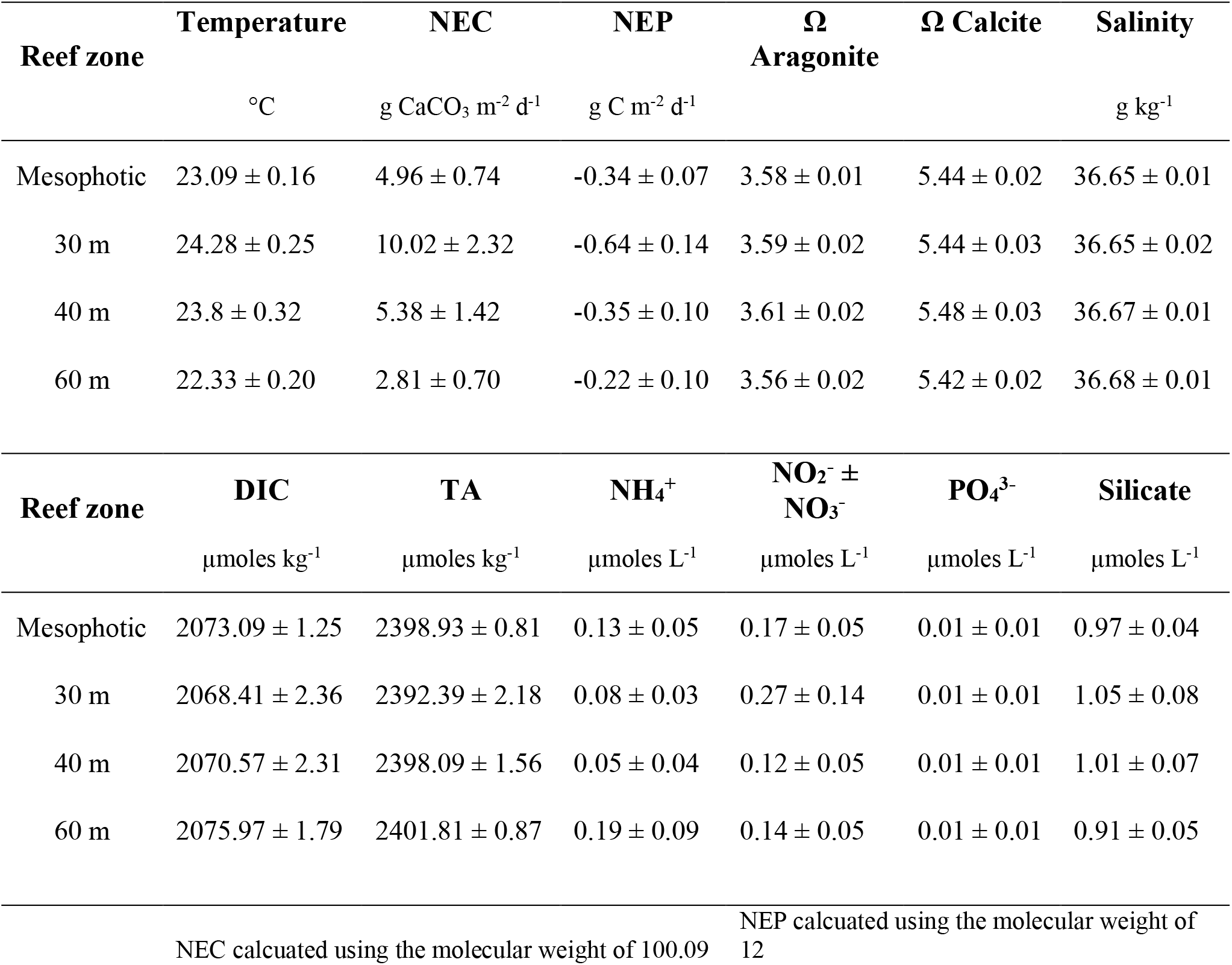
Summary of physico-chemical parameter averages ± SE over the duration of the study for the mesophotic coral reef and three reef zones. Note, nutrients were collected over the duration of the study at a subset of study locations.

| Reef zone | Temperature<br>°C | NEC<br>g CaCO<sub>3</sub> m<sup>-2</sup> d<sup>-1</sup> | NEP<br>g C m<sup>-2</sup> d<sup>-1</sup> | Ω<br>Aragonite | Ω Calcite | Salinity<br>g kg<sup>-1</sup> |
| --- | --- | --- | --- | --- | --- | --- |
| Mesophotic | 23.09 ± 0.16 | 4.96 ± 0.74 | -0.34 ± 0.07 | 3.58 ± 0.01 | 5.44 ± 0.02 | 36.65 ± 0.01 |
| 30 m | 24.28 ± 0.25 | 10.02 ± 2.32 | -0.64 ± 0.14 | 3.59 ± 0.02 | 5.44 ± 0.03 | 36.65 ± 0.02 |
| 40 m | 23.8 ± 0.32 | 5.38 ± 1.42 | -0.35 ± 0.10 | 3.61 ± 0.02 | 5.48 ± 0.03 | 36.67 ± 0.01 |
| 60 m | 22.33 ± 0.20 | 2.81 ± 0.70 | -0.22 ± 0.10 | 3.56 ± 0.02 | 5.42 ± 0.02 | 36.68 ± 0.01 |

| Reef zone | DIC<br>μmoles kg<sup>-1</sup> | TA<br>μmoles kg<sup>-1</sup> | NH<sub>4</sub><sup>+</sup><br>μmoles L<sup>-1</sup> | NO<sub>2</sub><sup>-</sup> ± NO<sub>3</sub><sup>-</sup><br>μmoles L<sup>-1</sup> | PO<sub>4</sub><sup>3-</sup><br>μmoles L<sup>-1</sup> | Silicate<br>μmoles L<sup>-1</sup> |
| --- | --- | --- | --- | --- | --- | --- |
| Mesophotic | 2073.09 ± 1.25 | 2398.93 ± 0.81 | 0.13 ± 0.05 | 0.17 ± 0.05 | 0.01 ± 0.01 | 0.97 ± 0.04 |
| 30 m | 2068.41 ± 2.36 | 2392.39 ± 2.18 | 0.08 ± 0.03 | 0.27 ± 0.14 | 0.01 ± 0.01 | 1.05 ± 0.08 |
| 40 m | 2070.57 ± 2.31 | 2398.09 ± 1.56 | 0.05 ± 0.04 | 0.12 ± 0.05 | 0.01 ± 0.01 | 1.01 ± 0.07 |
| 60 m | 2075.97 ± 1.79 | 2401.81 ± 0.87 | 0.19 ± 0.09 | 0.14 ± 0.05 | 0.01 ± 0.01 | 0.91 ± 0.05 |

|  |  |
| --- | --- |
| NEC calculated using the molecular weight of 100.09 | NEP calculated using the molecular weight of 12 |

### 3.2 Temporal changes in seawater carbonate chemistry and trophic status

The 2017 monthly observations of nDIC (Figure 2G-I) at the shallow mesophotic sites (30 m, 40 m) were generally higher than values calculated from BATS (∼3 - 4 μmoles kg^-1^). However, in 2018, the shallow reef observation fluctuated ± ∼ 1 μmoles kg^-1^ around those observed at BATS. Observations from the 2017 60 m reefs fluctuated ± ∼ 8 μmoles kg^-1^ around the BATS value whilst they were ∼ 8 μmoles kg^-1^ higher than values observed at BATS in 2018. The mean nDIC values (± standard deviation) for BATS, the 30 m, 40 m and 60 m MCEs over this study period were 2068.67 ± 10.90 μmoles kg^-1^, 2065.31 ± 10.37 μmoles kg^-1^, 2066.68 ± 11.20 μmoles kg^-1^ and 2071.46 ± 14.21 μmoles kg^-1^ respectively. Generally, the mesophotic reefs had lower nTA values than BATS for the duration of the study, with concentrations ranging between ∼ 2 – 45 μmoles kg^-1^ lower. The mean nTA values (Figure 2J-L) for BATS, the 30 m, 40 m, 60 m and MCEs over this study period were 2399.20 ± 10.90 μmoles kg^-1^, 2388.83 ± 12.94 μmoles kg^-1^, 2393.60 ± 7.53 μmoles kg^-1^ and 2396.62 ± 5.63 μmoles kg^-1^ respectively. The values recorded from mesophotic reefs followed the seasonal pattern recorded at BATS for both nDIC and nTA (Figure 2G-L). The saturation states for aragonite (Ω_aragonite_) and calcite (Ω_calcite_) remained stable for the duration of the study (Figure 2M-O).

With the exception of the 60 m reefs, the monthly mean NEP of MCEs exhibited similar albeit, inverted (i.e., decreases of NEP coincided with increases in NEC; Figure 3) seasonal patterns to those observed for NEC. There were seasonal differences in NEP signals recorded from the three reef zones (H = 17.144, *p* = <0.0001; Table 3) that reflected the trophic switch over the course of the year (Figure 3B,D,F). Net ecosystem production did not differ across the three reef zones (H = 3.728, *p* = 0.155) despite the observed differences in the monthly mean patterns. Between May and December, the 30 – 40 m NEP values were generally negative and symptomatic of autotrophy (photosynthesis > respiration; minimum value -3.14 g C m^-2^ d^-1^). Positive increases in NEP indicative with heterotrophy (i.e., photosynthesis < respiration) occurred later in the year and for longer time periods with increasing depth. Generally, periods of heterotrophy occurred in the winter for 30 m reefs, early summer in the 40 m reefs and throughout the summer and fall in the 60 m reefs (Figure 3B,D,F). Calcification (positive NEC; Figure 3A,C,E) generally occurred between May and December with the greatest rates measured at the 30 m depths and steadily decreased with increased depth (Z = 2.803, *p* = 0.015; Table 3). The peak monthly average (± standard deviation) calcification periods for all three depth ranges occurred in September (30 m = 36.62 ± 4.23 g CaCO_3_ m^-2^ d^-1^ in 2017, 40 m = 30.90 ± 0.01 g CaCO_3_ m^-2^ d^-1^ in 2017) and October (60 m = 16.44 ± 11.74 g CaCO_3_ m^-2^ d^-1^ in 2018).

**Figure 3.**
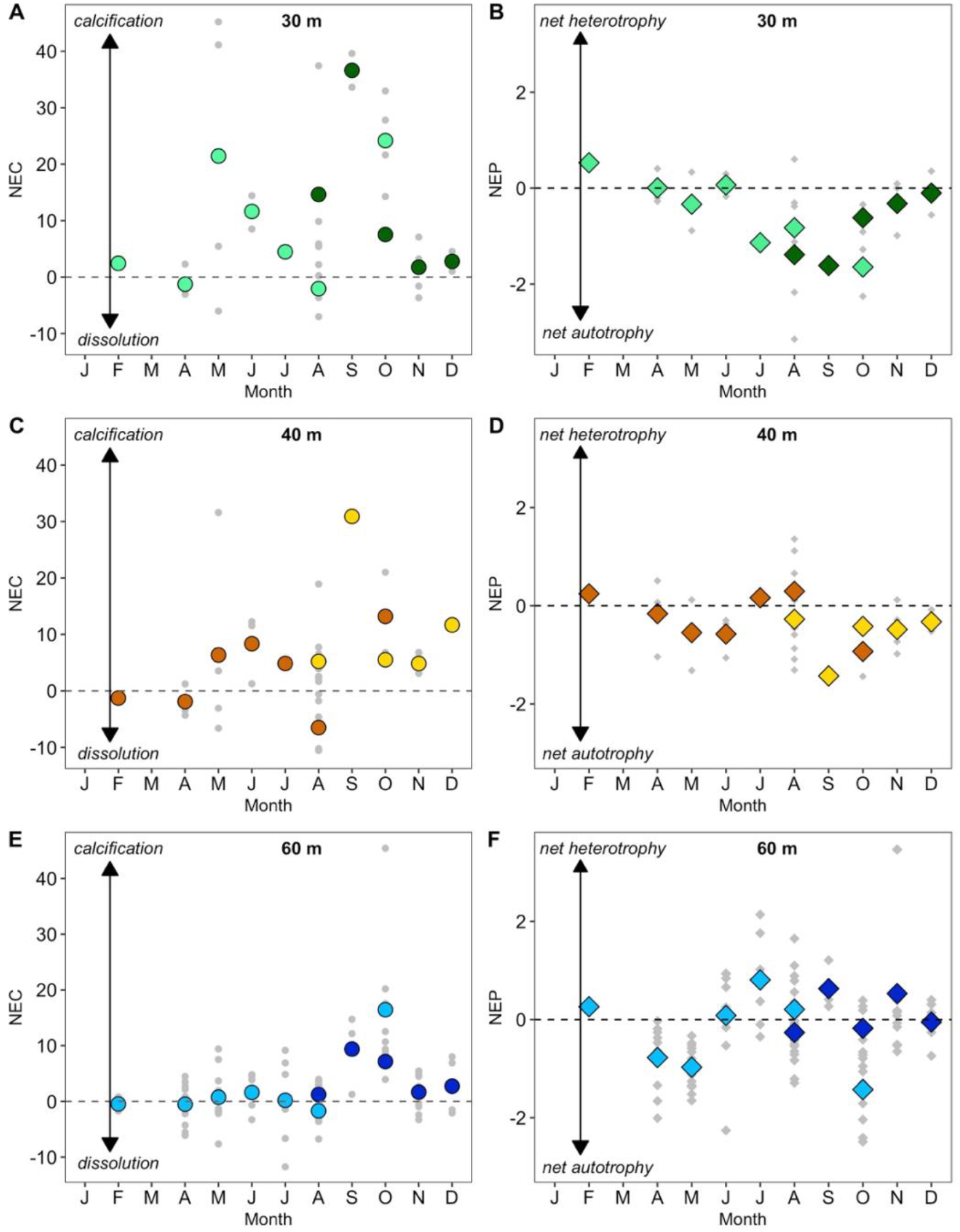
Mean seasonal climatology of net ecosystem calcification (NEC; g CaCO_3_ m^-2^ d^-1^) and net ecosystem production (NEP; g C m^-2^ d^-1^) for 30 m, 40m, 60 m reefs. Grey symbols represent actual samples **(A)** NEC, dark green circles denote monthly mean values for 2017, light green circles denote monthly mean values for 2018; **(B)** NEP, dark green diamonds denote monthly mean values for 2017, light green diamonds circles denote monthly mean values for 2018; **(C)** NEC, orange circles denote monthly mean values for 2017, dark orange circles denote monthly mean values for 2018; **(D)** NEP, orange diamonds denote monthly mean values for 2017, dark orange diamonds denote monthly mean values for 2018; **(E)** NEC, dark blue circles denote monthly mean values for 2017, light blue circles denote monthly mean values for 2018; **(F)** NEP, dark blue diamond’s denote monthly mean values for 2017, light blue diamond’s denote monthly mean values for 2018. Dashed lines equal calcification and trophic status are in balance (e.g., NEC = 0, NEP = 0).

**Table 3.** Summary of analyses statistically comparing Net Ecosystem Calcification (NEC) and Net Ecosystem Production (NEP) between sampling depth (m) and time of year (Kruskal-Wallis H test). Variation in community NEC and NEP between sampling depth (m) and time of year (Dunn’s test).

| NEC | Kruskal-Wallis H |  |  | NEC | Dunn's test |  |
| --- | --- | --- | --- | --- | --- | --- |
|  |  |  |  |  | (Kruskal-Wallis multiple comparison) |  |
|  | *df* | *H* | *P* |  | *Z* | *P.adj* |
| **Depth** | 2 | 9.013 | **0.011** | **Depth** |  |  |
|  |  |  |  | 30 m - 40 m | 0.862 | 0.388 |
|  |  |  |  | **30 m - 60 m** | 2.803 | **0.015** |
|  |  |  |  | 40 m - 60 m | 1.835 | 0.100 |
| **Season** | 3 | 35.686 | **<0.001** | **Season** |  |  |
|  |  |  |  | **Autumn - Spring** | 5.320 | **<0.001** |
|  |  |  |  | **Autumn - Summer** | 4.892 | **<0.001** |
|  |  |  |  | Spring - Summer | -1.213 | 0.270 |
|  |  |  |  | Autumn - Winter | 2.096 | 0.072 |
|  |  |  |  | Spring - Winter | -1.691 | 0.136 |
|  |  |  |  | Summer - Winter | -0.949 | 0.342 |
| NEP | Kruskal-Wallis H |  |  | NEP | Dunn's test |  |
|  |  |  |  |  | (Kruskal-Wallis multiple comparison) |  |
|  | *df* | *H* | *P* |  | *Z* | *P.adj* |
| **Depth** | 2 | 3.728 | 0.155 | **Depth** |  |  |
|  |  |  |  | 30 m - 40 m | -0.945 | 0.517 |
|  |  |  |  | 30 m - 60 m | -1.910 | 0.168 |
|  |  |  |  | 40 m - 60 m | -0.818 | 0.413 |
| **Season** | 3 | 17.144 | **<0.001** | **Season** |  |  |
|  |  |  |  | Autumn - Spring | 0.338 | 0.735 |
|  |  |  |  | **Autumn - Summer** | -2.931 | **0.010** |
|  |  |  |  | **Spring - Summer** | -2.939 | **0.020** |
|  |  |  |  | **Autumn - Winter** | -2.750 | **0.009** |
|  |  |  |  | **Spring - Winter** | -2.846 | **0.009** |
|  |  |  |  | Summer - Winter | -0.956 | 0.407 |

## 4 Discussion

### 4.1 Calcification Status

Bermudan mesophotic reefs exhibit both spatial and temporal variability in biogeochemical processes (i.e., the balance of photosynthesis, respiration, calcification, and CaCO_3_ dissolution). The mean NEC for the collective mesophotic reef system and individual reef depths investigated were positive thus indicative of net calcification (Figure 3A,C,E). The greatest rates of NEC were measured at the 30 m depths and steadily decreased with increased depth to the point that the 60 m reefs were in a state of equilibrium (calcification = dissolution) for ∼ 6 months of the year. The peak calcification period for all three depth ranges occurred in September and October followed by a period of equilibrium and or dissolution in the winter months. This switch between net accretion and net dissolution has been documented on a seasonal basis for Bermuda shallow reefs (Yeakel et al., 2015; Muehllehner et al., 2016; Bates, 2017; Cyronak et al., 2018). The peak calcification periods are comparable to a recent study on shallow reef calcification (Bates, 2017). The same study also recorded a reduction of accretion rates to near zero during an annual cycle over the duration of the 20-year time-series study. The monthly mean NEC (± standard deviation) for the 30 m, 40 m and 60 m reefs was 10.02 ± 14.32 g CaCO_3_ m^-2^ d^-1^, 5.38 ± 9.09 g CaCO_3_ m^-2^ d^-1^ and 2.81 ± 6.94 g CaCO_3_ m^-2^ d^-1^. These calcification rates are comparable to scaled *in situ* skeletal growth rates of the Grooved brain coral (*Diploria labryinthiformis* Linnaeus 1758; ∼1.30 -3.20 g CaCO_3_ m^-2^ d^-1^; Bates et al., 2010) located on the north coral platform of Bermuda at ∼10 m depth. However, these skeletal rates should not be taken as a direct comparison since the reef types and environmental conditions are not cognate (Andersson and Gledhill, 2013). Literature on mesophotic biogeochemistry and influences thereof are lacking (Hoegh-Guldberg et al., 2017), therefore it makes it impossible for direct NEC comparisons to other mesophotic locations but to give these values context, they fall within the range of average global coral reef NEC values 2.00 – 25.00 g CaCO_3_ m^-2^ d^-1^ (Atkinson, 2011). Interestingly, the 60 m locations show variability of calcification estimates (NEC). The three locations (Figure 1) spend differing accumulative time in a state of net calcification (BT1= 69%, BT2 = 76%, BT3 = 66%) over the duration of the study period (monthly; August 2017 – October 2018). The mean calcification rates for BT1, BT2 and BT3 were 2.63 ± 6.03g CaCO_3_ m^-2^ d^-1^, 7.38 ± 11.67 g CaCO_3_ m^-2^ d^-1^, and 4.40 ± 10.11 g CaCO_3_ m^-2^ d^-1^ respectfully. The reason for this variability is currently unknown but may be influenced by local hydrology. Bates (2017) determined that Bermuda shallow coral reef NEC have increased at approximately 3% per year (∼0.7 ± 0.3 g CaCO_3_ m^-2^ d^-1^) over a 20-year period (1996 – 2016). Since such mesophotic rate measurements constitute the first study, it is impossible to say if this trend will extend into mesophotic depths. However, both the study by Bates (2017) and a separate study by Yeakel et al. (2015) suggested episodic events of elevated NEC indicative of high calcification could be enhanced through alternative carbon sources (i.e., acquisition of organic nutrients through advection of biomass, e.g., zooplankton) as indicated by increased heterotrophy (> NEP; Figure 3E,F). This potential response appears to be evident in elevated measurements of NEP from the 60 m reefs in September and November 2017. Whilst autotrophy (photosynthesis by symbiont) is the primary energy source for most scleractinian corals, it has been demonstrated that up to ∼ 60% of the metabolic requirements of a coral (Houlbrèque and Ferrier-Pagès, 2009) can be supplied through heterotrophy (i.e., incorporation of particulate and dissolved organic matter, respiration). Mesocosm based feeding experiments have shown this input of organic carbon can maintain calcification rates under ocean acidification conditions (Drenkard et al., 2013; Towle et al., 2015) as well as enhance photosynthesis (Houlbrèque and Ferrier-Pagès, 2009). Yeakel et al. (2015) postulate these changes in metabolite source led to elevated summertime calcification rates (NEC), draw down of nTA and a reduction in pH and Ω_aragonite_. These high calcification / acidification events are correlated with a negative winter North Atlantic Oscillation (NAO). One could hypothesize that through geographical location, mesophotic reefs, at least for Bermuda, are the “boundary layer” between the open ocean and shallow reefs. One would surmise that any benefits episodic events such as the winter NAO afford shallow corals through advection of biomass onto the reef, could be happening on a more frequent basis for mesophotic reefs. A study of coral trophic zonation on Palmyra Atoll determined that internal waves (example of a transport mechanism of oceanic plankton) were depth restricted with only 4 – 8 % of events extending up reef slopes shallower than 30 m (Williams et al., 2018). Increases in both phytoplankton and zooplankton biomass have been documented at the Bermuda Atlantic Time-series Study site (∼ 80 km southeast of Bermuda), with resultant increases in active carbon flux due to diel vertical migration (zooplankton) and passive carbon flux by faecal pellets export (i.e., particulate organic matter; Steinberg et al., 2012).

The three locations were all accumulatively in a state of net heterotrophy for ∼ 30 % of the study period, however BT3 had the greatest accumulative time at 38%. Whilst elevated calcification levels and corresponding draw down of nTA were observed on the 60 m reefs in the latter part of 2017 (September and November), there appeared to be no reduction in Ω_aragonite_ or pH. Whilst there were increases in the heterotrophy signal on the 60 m reef during this period, the same responses were not observed on the 30 m or 40 m reefs. In fact, there were stronger periods of mean autotrophy during September relative to the previous month. This trend also occurred in November on the 40 m reefs. It would be expected that there would be an increased reliance on heterotrophy with increased depth by scleractinian corals (Williams et al., 2018) due to increased light attenuation and increased particulate resource availability (Fox et al., 2018) driven by hydrodynamic processes such as upwelling and internal waves.

The reason(s) for the apparent variations in trophic status between the three reef zones is currently unknown however, it is suspected that fine-scale hydrodynamic and hydrographic regimes are likely drivers of these spatial disparities (Williams et al., 2018). Longer term measurements of mesophotic biogeochemistry and a better understanding of hydrology will (1) validate the theory of augmented calcification through increased nutrition (i.e., heterotrophy); (2) help delineate the environmental controls on these deeper reef systems.

In addition to the advection of biomass (e.g., plankton and POM) from oceanic sources, deep-water upwelling and internal waves are known to influence nutrient availability through influxes of inorganic nutrients onto reef systems (Stuhldreier et al., 2015). These allochthonous inputs are often rapidly converted to particulate resources therefore leading to increased primary productivity. The increase in POM resource availability enables coral heterotrophy and is a critical component of fish productivity which can be sustained through multiple heterotrophy trophic pathways (Chassot et al., 2010; Morais and Bellwood, 2019).

Primary productivity within coral reef systems has traditionally been viewed as the regulator of higher trophic levels (bottom-up control). However, it has been postulated that top-down control (biomass altered by predation) enables fish communities to influence biogeochemical cycling rates (Kavanagh and Galbraith, 2018), for example through consumer-mediated nutrient dynamics (Allgeier et al., 2017). Primary production would potentially be enhanced through the excretion and egestion of essential nutrients.

### 4.2 Trophic Status

The three reef systems exhibited differences in estimated trophic status over the course of the study. The monthly mean NEP estimates for the 30 m reefs between February and June were generally negative and symptomatic of autotrophy (photosynthesis > respiration). During this same timeframe, the 40 m reef systems switched to a state of net autotrophy during May and June. The 60 m reefs were generally in a state of equilibrium or net autotrophy during this period followed by positive NEP indicative with heterotrophy (i.e., photosynthesis < respiration) or equilibrium between June – February. This pattern of net autotrophy in the early summer with a shift to strong heterotrophy in late summer was described by Bates et al. (2010) as the “Carbonate Chemistry Coral Reef Ecosystem Feedback” (CREF hypothesis). During the summer months, elevated autotrophy (e.g., scleractinian coral calcifying) heighten the Ω_aragonite_, and [CO_3_^2-^] conditions (e.g., CO_2_ uptake and photosynthesis). In the late summer, there is a switch in metabolic source, CO_2_ released through respiration leads to a suppression of photosynthetic activity. Evidence for this can be seen in Figure 3B,D,F albeit the estimated NEP values do not become positive (i.e., heterotrophic) for either the 30 m or 40 m reefs. Instead, the feedback causes a strong reduction in autotrophy closer to a state of equilibrium. Whilst there was variability between the three reefs depths, the overall status of the mesophotic system was net autotrophic (−0.34 ± 0.92 g C m^-2^ d^-1^) and not in a state of balance. This determination is the opposite of the trophic evaluation for Bermuda shallow reefs (net heterotrophic; + 0.20 ± 0.9 g C m^-2^ d^-1^). These findings present an interesting conundrum. Zooxanthellate corals are generally restricted to depths where light levels typically exceed the 0.5% of the subsurface intensity (Dubinsky and Stambler, 2011). Therefore, light availability is a primary factor that drives the vertical zonation of communities. The classical viewpoint would be to expect that the shallow water reefs would be more likely to derive carbon by way of the inorganic carbon cycle (i.e., photosynthesis) due to greater levels of surface irradiance. However, as discussed in the calcification section, there are energetic benefits when zooxanthellate corals utilize both inorganic and organic carbon sources (photosynthesis + respiration). It does raise the question about the exact composition of the primary calcifiers at mesophotic depths and how representative are these rate measurements and if we truly are “taking the metabolic pulse” of these communities (Cyronak et al., 2018). To further complicate our understanding of these complex biogeochemical processes, there are alternative inputs of CaCO_3_ and alkalinity fluxes that have not been considered. Scleractinian corals are considered dominant calcifiers on reef systems, however all marine teleosts produce and excrete CaCO_3_ as an osmoregulatory product due to the constant swallowing of seawater (Wilson et al., 2009). Calcium carbonate precipitates into the digestive tract and is excreted either as pellets or with faecal matter which is estimated to contribute ∼ 3 – 15% of total new CaCO_3_ production to the upper oceanic environment (Wilson et al., 2009). Dissolution of the excreted CaCO_3_ would lead to increases in total alkalinity. It should be noted that the calculated NEC rates are a relative expression of the balance of calcification and dissolution based on observed differences between offshore and *in situ* normalised TA measurements (Equation 4). Increases in offshore TA values would result in a stronger positive NEC signal. Alternatively, increases in mesophotic TA values would correspond to a stronger negative NEC signal. Hypothetically, fluctuations in fish abundances at either survey or reference sites (e.g., diel vertical migration of deep-sea fish) could lead to direct changes in biogeochemical processes and alternative interpretation of calcification/dissolution results.

### 4.3 Other considerations

Knowledge on benthic community composition for mesophotic reefs is lacking on a global scale (Loya et al., 2019). Goodbody-Gringley et al. (2019) summarise known information for Bermuda and a recent field guide to Bermuda’s MCEs has been produced (Stefanoudis et al., 2018). Based on a limited number of quantitative surveys (n = 5) algae and a sand/rubble complex account for 69% and ∼25% of the broad functional groupings of the benthic community at 60 m depths. For context, Scleractinia represent 0.02% of these data. It could be postulated that these measurements of calcification/dissolution and autotrophic/heterotrophic may not be directly coupled to coral calcification and may in fact be an alternative signal such as a seasonal reduction in CaCO_3_ dissolution/bioerosion or fish movement patterns. Within the algal grouping, encrusting crustose coralline algae (CCA) and rhodoliths are the main calcifying taxa. Rhodoliths and CCA perform valuable ecosystem services of substrate provision through calcification. Which carbonate mineral phase (i.e., aragonite, calcite, and magnesian calcite - Mg-calcite) these red algae utilize for calcification is taxa specific (Nash et al., 2019) but in the case of *Corallinales*, the carbonate mineral phase is Mg-calcite (Nash et al., 2011). Gorgonians are known to calcify using Mg-calcite and are one of the most diverse coral groups on Caribbean MCES below 60 m depths. A recent study on the effects of ocean acidification on *Corallium rubrum* (Linnaeus, 1758) demonstrated lower pH (7.81 pH) significantly reduced skeletal growth (Bramanti et al., 2013), therefore till disproven, one could assume a similar response by mesophotic *Corallinales*. The effect of ocean acidification (OA) and the reduction of seawater saturation state on marine organisms’ ability to accrete CaCO_3_ has been well documented in the literature. However, studies do not tend to delineate between different carbonate mineral phase (i.e., aragonite, calcite, and Mg-calcite) saturation states (Lebrato et al., 2016) often referring to fluctuations in Ω_aragonite_ or Ω_calcite_. Magnesium calcite minerals that have a magnesium content greater than 8 – 12 mol% are more soluble than both aragonite and calcite (Gattuso and Hansson, 2011). New evidence suggests Ω_aragonite_ or Ω_calcite_ do not account for the Mg content of calcite (increased solubility) therefore are not appropriate estimates of seawater saturation state with respect to Mg-calcite (Lebrato et al., 2016). The same study determined that 24% of benthic calcite producing calcifiers are currently experiencing under saturated conditions (i.e., dissolution; Ω_Mgcalcite-x_). Of those, the majority (95%) were found in the tropics.

A lack of long-term measurements currently restricts our ability to interpret natural and seasonal variability of mesophotic biogeochemistry. Ultimately, this hinders our capacity to predict biogeochemical responses of these environments to future ocean acidification and climate change scenarios. The predicted effects of OA on calcifying benthic communities are often based on the relationship between average Ω_aragonite_ and NEC (gross calcification – gross CaCO3 dissolution) which typically changes at a rate of 102% NEC per unit change of seawater Ω_aragonite_ (Eyre et al., 2018). Predicting OA driven changes to mesophotic coral ecosystems is beyond the scope of this study. However, the estimates of NEC and NEP derived by this investigation represent the first known biogeochemical measurements for mesophotic reefs and therefore provided the critical first step towards enabling our predictive capabilities. The unique location of the study allows these measurements to be considered in the context of contemporaneous offshore changes observed at the Bermuda Atlantic Time-series Study (BATS) site. The BATS program was established in 1988 and represents the longest-running time-series for biogeochemical oceanographic data.

It should be recognized that these measurements are of the balance between calcification and CaCO_3_ dissolution (NEC) and organic carbon cycling (NEP). As such, they might not be directly coupled to benthic calcification (i.e., estimates of framework / sediment transport) or represent specific energy pathways (e.g., CaCO_3_ sediment dissolution = increase in DIC). However, the NEC estimates establish that the seawater chemistry of Bermuda mesophotic reefs is chemically conducive for calcification. These reef systems are in a net state of calcification (i.e., accretion of CaCO_3_) and exhibited changes in calcification (strongest periods in the late summer) and trophic state (switch heterotrophy to autotrophy) during the investigation. These findings represent a unique yet limited snapshot of the biogeochemical conditions for Bermuda’s mesophotic reefs. Decoupling the environmental controls exerted on these mesophotic reefs in general, will require considerably longer observational time scales coupled with experimental approaches.

During the 18-month time frame of this study, mesophotic biogeochemical observations displayed a similar response to the seasonal trends established for adjacent shallow reefs (Yeakel et al., 2015; Bates, 2017). All three mesophotic reef zones were net accretive (i.e., gross calcification > gross CaCO3 dissolution) and in a net state of autotrophy. Despite this, these systems exhibited periods of variability through a trophic switch between autotrophy and heterotrophy. It remains to be determined if these signals are indicative of the seasonal variability established on Bermuda’s shallow water reef system.

Whilst these data are invaluable and have begun to fill much needed knowledge gaps, the study has generated additional questions that require further examination. For example, why are the mesophotic reefs in opposing trophic states to those of the shallow reef counter parts (MCE = net autotrophic, shallow reefs = net heterotrophic; Bates, 2017)? The values established by this study demonstrate just how close these understudied ecosystems are in terms of the known boundary thresholds for low saturation states of reefs (Figure 2M-O). Making predictions on how these ecosystems will respond to future climate changes will be extremely difficult when it is not currently known if the biogeochemical signals are a true representation of the status of these reefs, or alternatively signals of seasonal reduction in CaCO_3_ dissolution/bioerosion. Although these processes are yet to be fully understood, the apparent increase in NEC and NEP rates (∼30%) on Bermudan shallow reefs over the last 20 years (Bates, 2017) are an indication that there is a level of resilience to the changing environment that may extend to MCEs. At what level and for how long are critical questions that will need to be urgently addressed in the near future.

## Conflict of Interest

The authors declare that the research was conducted in the absence of any commercial or financial relationships that could be construed as a potential conflict of interest.

## Author Contributions

TJN conceive the idea and designed the study, analyzed the results, prepared the figures and wrote the manuscript. RG performed seawater analyzes and NRB provided comparative data for various components of the study. TN wrote the first draft of the manuscript, RB and NRB contributed to manuscript revision, read, and approved the submitted version.

## Funding

This research was produced with financial support from the European Union through the BEST2.0+ Program (#1634, #2274), BIOS Bermuda Program, BIOS Grant-in-Aid program, BIOS UK Associates partnership, and the National Science Foundation’s Diversity of Ocean Sciences: Research Experience for Undergraduates (NSF #1757475).

## Acknowledgments

Except for comparative data utilized from the Bermuda Atlantic Time-series Study, all data presented in this manuscript is the product of Timothy Noyes PhD thesis ‘Determining the spatial and temporal trends of mesophotic fish biodiversity and reef-scale calcification using novel approaches’, Manchester, University of Salford. The following are thanked for their contribution to sampling efforts, Rosie Dowell, Jonas Schroder, Emma O’Donnell, Ellie Corbet and Alex Lundberg. Kaitlin Noyes, Christopher Noyes, and Stefano Mariani provided comments that improved an earlier version of this manuscript.

## Notes

### Competing Interest Statement

The authors have declared no competing interest.

## References

Allgeier, J. E., Burkepile, D. E., and Layman, C. A. (2017). Animal pee in the sea: consumer-mediated nutrient dynamics in the world’s changing oceans. Glob Chang Biol 23, 2166–2178. doi: 10.1111/gcb.13625.

Andersson, A. J., and Gledhill, D. (2013). Ocean Acidification and Coral Reefs: Effects on Breakdown, Dissolution, and Net Ecosystem Calcification. Ann Rev Mar Sci 5, 321–348. doi: 10.1146/annurev-marine-121211-172241.

Andersson, A. J., Kuffner, I. B., MacKenzie, F. T., Jokiel, P. L., Rodgers, K. S., and Tan, A. (2009). Net Loss of CaCO3 from a subtropical calcifying community due to seawater acidification: Mesocosm-scale experimental evidence. Biogeosciences 6, 1811–1823. doi: 10.5194/bg-6-1811-2009.

Andersson, A. J., Yeakel, K. L., Bates, N. R., and De Putron, S. J. (2014). Partial offsets in ocean acidification from changing coral reef biogeochemistry. Nat Clim Chang 4, 56–61. doi: 10.1038/nclimate2050.

Atkinson, M. J. (2011). “Biogeochemistry of nutrients,” in Coral reefs: An ecosystem in transition, eds. Z. Dubinsky and S. Stambler (Springer Netherlands), 199–206. doi: doi.org/10.1007/978-94-007-0114-4_13.

Baker, E., Puglise, K., and Harris, P. (2016). Mesophotic Coral Ecosystems -A Lifeboat for Coral Reefs? The United Nations Environment Programme and GRID-Arendal, Nairobi and Arendal.

Bates, N. R. (2002). Seasonal variability of the effect of coral reefs on seawater CO 2 and air-sea CO 2 exchange.

Bates, N. R. (2017). Twenty years of marine carbon cycle observations at Devils Hole Bermuda provide insights into seasonal hypoxia, coral reef calcification, and ocean acidification. Front Mar Sci 4, 1–23. doi: 10.3389/fmars.2017.00036.

Bates, N. R., Amat, A., and Andersson, A. J. (2010). Feedbacks and responses of coral calcification on the Bermuda reef system to seasonal changes in biological processes and ocean acidification. Biogeosciences 7, 2509–2530. doi: 10.5194/bg-7-2509-2010.

Bates, N. R., Best, M. H. P., Neely, K., Garley, R., Dickson, A. G., and Johnson, R. J. (2012). Detecting anthropogenic carbon dioxide uptake and ocean acidification in the North Atlantic Ocean. Biogeosciences 9, 2509–2522. doi: 10.5194/bg-9-2509-2012.

Bates, N. R., Michaels, A. F., and Knap, A. H. (1996). Alkalinity changes in the Sargasso Sea: of calcification? geochemical evidence.

Bongaerts, P., Muir, P., Englebert, N., Bridge, T. C. L., and Hoegh-Guldberg, O. (2013). Cyclone damage at mesophotic depths on Myrmidon Reef (GBR). Coral Reefs 32, 935. doi: 10.1007/s00338-013-1052-y.

Bongaerts, P., Ridgway, T., Sampayo, E. M., and Hoegh-Guldberg, O. (2010). Assessing the “deep reef refugia” hypothesis: Focus on Caribbean reefs. Coral Reefs 29, 1–19. doi: 10.1007/s00338-009-0581-x.

Bongaerts, P., Riginos, C., Brunner, R., Englebert, N., Smith, S. R., and Hoegh-Guldberg, O. (2017). Deep reefs are not universal refuges: Reseeding potential varies among coral species. Sci Adv 3. doi: 10.1126/sciadv.1602373.

Bridge, T. C. L., Hughes, T. P., Guinotte, J. M., and Bongaerts, P. (2013). Call to protect all coral reefs. Nat Clim Chang 3, 528–530. doi: 10.1038/nclimate1879.

Chassot, E., Bonhommeau, S., Dulvy, N. K., Mélin, F., Watson, R., Gascuel, D., et al. (2010). Global marine primary production constrains fisheries catches. Ecol Lett 13, 495–505. doi: 10.1111/j.1461-0248.2010.01443.x.

Cinner, J. E., Huchery, C., MacNeil, M. A., Graham, N. A. J., McClanahan, T. R., Maina, J., et al. (2016). Bright spots among the world’s coral reefs. Nature 535, 416–419. doi: 10.1038/nature18607.

Coates, K. A., Fourqurean, J. W., Kenworthy, W. J., Logan, A., Manuel, S. A., and Smith, S. R. (2013). “Introduction to Bermuda: Geology, Oceanography and Climate,” in, 115–133. doi: 10.1007/978-94-007-5965-7_10.

Cohen, A. (2003). Geochemical Perspectives on Coral Mineralization. Rev Mineral Geochem 54, 151–187. doi: 10.2113/0540151.

Colin, P. L. (2009). Marine environments of Palau. San Diego: Indo-Pacific Press.

Colin, P. L., and Lindfield, S. J. (2019). “Palau,” in Mesophotic Coral Ecosystems Coral Reefs of the World 12, eds. Y. Loya, K. Puglise, and T. C. L. Bridge (Springer International Publishing), 285–320. doi: 10.1007/978-3-319-9275-0_16.

Courtney, T. A., Andersson, A. J., Bates, N. R., Collins, A., Cyronak, T., de Putron, S. J., et al. (2016). Comparing chemistry and census-based estimates of net ecosystem calcification on a rim reef in Bermuda. Front Mar Sci 3. doi: 10.3389/fmars.2016.00181.

Courtney, T. A., Cyronak, T., Griffin, A. J., and Andersson, A. J. (2021). Implications of salinity normalization of seawater total alkalinity in coral reef metabolism studies. PLoS One 16, 1–13. doi: 10.1371/journal.pone.0261210.

Cyronak, T., Andersson, A. J., Langdon, C., Albright, R., Bates, N. R., Caldeira, K., et al. (2018). Taking the metabolic pulse of the world’s coral reefs. PLoS One 13. doi: 10.1371/journal.pone.0190872.

Dickson, A. G., and Millero, F. J. (1987). A comparison of the equilibrium constants for the dissociation of carbonic acid in seawater media. Deep Sea Research Part A, Oceanographic Research Papers 34, 1733–1743. doi: 10.1016/0198-0149(87)90021-5.

Dickson, A. G., Sabine, C. L., and Christian, J. R. (2007). Guide to Best Practices for Ocean CO2 measurements. North Pacific Marine Science Organization.

Dove, S. G., Kline, D. I., Pantos, O., Angly, F. E., Tyson, G. W., and Hoegh-Guldberg, O. (2013). Future reef decalcification under a business-as-usual CO2 emission scenario. Proc Natl Acad Sci U S A 110, 15342–15347. doi: 10.1073/pnas.1302701110.

Drenkard, E. J., Cohen, A. L., McCorkle, D. C., de Putron, S. J., Starczak, V. R., and Zicht, A. E. (2013). Calcification by juvenile corals under heterotrophy and elevated CO2. Coral Reefs 32, 727–735. doi: 10.1007/s00338-013-1021-5.

Dubinsky, Z., and Stambler, N. (2011). Coral reefs: An ecosystem in transition. Coral Reefs: An Ecosystem in Transition, 1–552. doi: 10.1007/978-94-007-0114-4.

Eyre, B. D., Cyronak, T., Drupp, P., De Carlo, E. H., Sachs, J. P., and Andersson, A. J. (2018). Coral reefs will transition to net dissolving before end of century. Science (1979) 359, 908–911. doi: 10.1126/science.aao1118.

Fox, M. D., Williams, G. J., Johnson, M. D., Radice, V. Z., Zgliczynski, B. J., Kelly, E. L. A., et al. (2018). Gradients in Primary Production Predict Trophic Strategies of Mixotrophic Corals across Spatial Scales. Current Biology 28, 3355–3363.e4. doi: 10.1016/j.cub.2018.08.057.

Fricke, H., and Meischner, D. (1985). Depth limits of Bermudan scleractinian corals: a submersible survey.

Gattuso, Jean-Pierre, and Lina Hansson (eds), Ocean Acidification (Oxford, 2011, Oxford Academic, doi: 10.1093.

Glynn, P. W. (1996). Coral reef bleaching: Facts, hypotheses and implications. Glob Chang Biol 2, 495–509. doi: 10.1111/j.1365-2486.1996.tb00063.x.

Goodbody-Gringley, G., Marchini, C., Chequer, A. D., and Goffredo, S. (2015). Population structure of Montastraea cavernosa on shallow versus mesophotic reefs in Bermuda. PLoS One 10. doi: 10.1371/journal.pone.0142427.

Goodbody-Gringley, G., Noyes, T., and Smith, S. R. (2019). “Bermuda,” in Mesophotic Coral Ecosystems, Coral Reefs of the World 12, eds. L. Yossi, K. A. Puglise, and T. Bridge (Springer International Publishing), 31–45. doi: 10.1007/978-3-319-92735-0_2.

Hinderstein, L. M., Marr, J. C. A., Martinez, F. A., Dowgiallo, M. J., Puglise, K. A., Pyle, R. L., et al. (2010). Theme section on “Mesophotic Coral Ecosystems: Characterization, Ecology, and Management.” Coral Reefs 29, 247–251. doi: 10.1007/s00338-010-0614-5.

Hoegh-Guldberg, O. (2011). Coral reef ecosystems and anthropogenic climate change. Reg Environ Change 11, 215–227. doi: 10.1007/s10113-010-0189-2.

Hoegh-Guldberg, O., Mumby, P. J., Hooten, A. J., Steneck, R. S., Greenfield, P., Gomez, E., et al. (2007). Coral reefs under rapid climate change and ocean acidification. Science (1979) 318, 1737–1742. doi: 10.1126/science.1152509.

Hoegh-Guldberg, O., Poloczanska, E. S., Skirving, W., and Dove, S. (2017). Coral reef ecosystems under climate change and ocean acidification. Front Mar Sci 4. doi: 10.3389/fmars.2017.00158.

Houlbrèque, F., and Ferrier-Pagès, C. (2009). Heterotrophy in Tropical Scleractinian Corals. Biological Reviews 84, 1–17. doi: 10.1111/j.1469-185X.2008.00058.x.

Hughes, T. P., Rodrigues, M. J., Bellwood, D. R., Ceccarelli, D., Hoegh-Guldberg, O., McCook, L., et al. (2007). Phase Shifts, Herbivory, and the Resilience of Coral Reefs to Climate Change. Current Biology 17, 360–365. doi: 10.1016/j.cub.2006.12.049.

Jackson, J. B. C., Kirby, M. X., Berger, W. H., Bjorndal, K. A., Botsford, L. W., Bourque, B. J., et al. (2001). Historical overfishing and the recent collapse of coastal ecosystems. Science (1979) 293, 629–637. doi: 10.1126/science.1059199.

Jackson, J., Donovan, M., Cramer, K., and Lam, V. eds. (2014). Status and Trends of Caribbean Coral Reefs : 1970-2012. Gland, Switzerland: Global Coral Reef Monitoring Network, IUCN doi: 10.1016/0377-8401(86)90099-4.

Jones, R., Johnson, R., Noyes, T., and Parsons, R. (2012). Spatial and temporal patterns of coral black band disease in relation to a major sewage outfall. Mar Ecol Prog Ser 462, 79–92. doi: 10.3354/meps09815.

Kavanagh, L., and Galbraith, E. (2018). Links between fish abundance and ocean biogeochemistry as recorded in marine sediments. PLoS One 13. doi: 10.1371/journal.pone.0199420.

Kleypas, J. A., Buddemeier, R. W., Archer, D., Gattuso, J. P., Langdon, C., and Opdyke, B. N. (1999). Geochemical consequences of increased atmospheric carbon dioxide on coral reefs. Science (1979) 284, 118–120. doi: 10.1126/science.284.5411.118.

Kleypas, J. A., Buddemeier, R. W., and Gattuso, J. P. (2001). The future of Coral reefs in an age of global change. International Journal of Earth Sciences 90, 426–437. doi: 10.1007/s005310000125.

Knap, A., Michaels, A. F., Steinberg, D. K. D., Bahr, F., Bates, N., Bell, S., et al. (1997). BATS Methods Manual. Version 4. U.S. JGOFS Planning Office, Woods Hole.

Langdon, C., Gattuso, J.-P., and Andersson, A. (2010). “Measurements of calcification and dissolution of benthic organisms and communities,” in Guide to beast practices for ocean acidification research and data reporting, eds. U. Riebesell, V. Fabry, L. Hansson, and J.-P. Gattuso (Luxembourg: Publications of the European Union), 213–232.

Langdon, C., Takahashi, T., Sweeney, C., Chipman, D., and Atkinson, J. (2000). rate of an experimental coral reef responds to manipulations in the concentrations of both Ca CO •. Global Biogeochem Cycles 14, 639–654.

Lantz, C. A., Atkinson, M. J., Winn, C. W., and Kahng, S. E. (2014). Dissolved inorganic carbon and total alkalinity of a Hawaiian fringing reef: Chemical techniques for monitoring the effects of ocean acidification on coral reefs. Coral Reefs 33, 105–115. doi: 10.1007/s00338-013-1082-5.

Lebrato, M., Andersson, A., Ries, J., Aronson, R., Lamare, M., Koeve, W., et al. (2016). Benthic marine calcifiers coexist with CaCO3-undersaturated seawater worldwide. Global Biogeochem Cycles 30, 1–16. doi: 10.1002/2015GB005260.Received.

Lewis, E., and Wallace, D. (1998). Program developed for CO2 system calculations. Ornl/Cdiac-105, 1–21. doi: 4735.

Logan, A. (1988). The Holocene Reefs of Bermuda. Sedimenta XI, 63.

Logan, A., and Murdoch, T. (2011). “Bermuda,” in Encyclopedia of modern coral reefs: structure, form and process, Earth Science Series., ed. D. Hopley (Dordrecht: Springer-Verlag), 469–486. doi: 10.1007/978-90-481-2639-2_46.

Loya, Y., Eyal, G., Treibitz, T., Lesser, M. P., and Appeldoorn, R. (2016). Theme section on mesophotic coral ecosystems: advances in knowledge and future perspectives. Coral Reefs 35, 1–9. doi: 10.1007/s00338-016-1410-7.

Loya, Y., Pulglise, K. A., and Bridge, T. C. L. eds. (2019). Mesophotic Coral Ecosystems Coral Reefs of the World 12. doi: 10.1007/978-3-319-92735-0.

Mehrbach, C., Culberson, C. H., Hawley, J. E., and Pytkowicx, R. M. (1973). Measurement Of The Apparent Dissociation Constants Of Carbonic Acid In Seawater At Atmospheric Pressure. Limnol Oceanogr 18, 897–907. doi: 10.4319/lo.1973.18.6.0897.

Moberg, F., and Folke, C. (1999). Ecological goods and services of coral reef ecosystems. Ecological Economics 29, 215–233. doi: 10.1016/S0921-8009(99)00009-9.

Morais, R. A., and Bellwood, D. R. (2019). Pelagic Subsidies Underpin Fish Productivity on a Degraded Coral Reef. Current Biology 29, 1521–1527.e6. doi: 10.1016/j.cub.2019.03.044.

Muehllehner, N., Langdon, C., Venti, A., and Kadko, D. (2016). Dynamics of carbonate chemistry, production, and calcification of the Florida Reef Tract (2009–2010): Evidence for seasonal dissolution. AGU Publications 30, 661–688. doi: 10.1002/2015GB005327.Received.

Nash, M. C., Diaz-Pulido, G., Harvey, A. S., and Adey, W. (2019). Coralline algal calcification: A morphological and process-based understanding. doi: 10.1371/journal.pone.0221396.

Nash, M. C., Troitzsch, U., Opdyke, B. N., Trafford, J. M., Russell, B. D., and Kline, D. I. (2011). First discovery of dolomite and magnesite in living coralline algae and its geobiological implications. Biogeosciences 8, 3331–3340. doi: 10.5194/bg-8-3331-2011.

Noyes, T. (2023) Determining the spatial and temporal trends of mesophotic fish biodiversity and reef-scale calcification using novel approaches. PhD. Manchester, University of Salford.

Orr, J. C., Fabry, V. J., Aumont, O., Bopp, L., Doney, S. C., Feely, R. A., et al. (2005). Anthropogenic ocean acidification over the twenty-first century and its impact on calcifying organisms. Nature 437, 681–686. doi: 10.1038/nature04095.

Pandolfi, J. M., Bradbury, R. H., Sala, E., Hughes, T. P., Bjorndal, K. A., Cooke, R. G., et al. (2003). Global trajectories of the long-term decline of coral reef ecosystems. Science (1979) 301, 955–958. doi: 10.1126/science.1085706.

Pandolfi, J. M., Connolly, S. R., Marshall, D. J., and Cohen, A. L. (2011). Projecting coral reef futures under global warming and ocean acidification. Science (1979) 333, 418–422. doi: 10.1126/science.1204794.

Puglise, K., Hinderstein, L., Marr, J., Dowgiallo, M., and Martinez, F. (2008). Mesophotic Coral Ecosystems Research Strategy. Silver Spring.

Pyle, R. L., Copus, J. M., Pyles, R. L., and Copus, J. M. (2019). “Mesophotic Coral Ecosystems: Introduction and Overview,” in Mesophotic Coral Ecosystems, Coral Reefs of the World 12, eds. Y. Loya, K. A. Puglise, and T. Bridge (Springer International Publishing), 3–27. doi: 10.1007/978-3-319-92735-0.

Romanó de Orte, M., Koweek, D. A., Cyronak, T., Takeshita, Y., Griffin, A., Wolfe, K., et al. (2021). Unexpected role of communities colonizing dead coral substrate in the calcification of coral reefs. Limnol Oceanogr 66, 1793–1803. doi: 10.1002/lno.11722.

Sarkis, S., van Beukering, P. J. H., McKenzie, E., Brander, L., Hess, S., Bervoets, T., et al. (2013). “Total Economic Value of Bermuda’s Coral Reefs: A Summary,” in Coral Reefs of the United Kingdom Overseas Territories, ed. C.R.C. Sheppard (Dordrecht: Springer Science and Business Media LLC), 201–211. doi: 10.1007/978-94-007-5965-7_15.

Semmler, R. F., Hoot, W. C., and Reaka, M. L. (2017). Are mesophotic coral ecosystems distinct communities and can they serve as refugia for shallow reefs? Coral Reefs 36, 433–444. doi: 10.1007/s00338-016-1530-0.

Smith, S. v., and Key, G. S. (1975). Carbon dioxide and metabolism in marine environments. Limnol Oceanogr 20, 493–495. doi: 10.4319/lo.1975.20.3.0493.

Spalding, M., Ravilious, C., and Green, E. (2001). World atlas of coral reefs. doi: 10.5860/choice.39-2540.

Stefanoudis, P. V., Rivers, M., Smith, S. R., Schneider, C. W., Wagner, D., Ford, H., et al. (2019). Low connectivity between shallow, mesophotic and rariphotic zone benthos. R Soc Open Sci 6, 190958. doi: 10.1098/rsos.190958.

Stefanoudis, P. v, Smith, S. R., Schneider, C., Wagner, D., Goodbody-Gringley, G., Xavier, J., et al. (2018). Bermuda Benthic Marine Life Field Identification Guide Bermuda Benthic Marine Life Field Identification Guide Deep Reef Benthos of Bermuda: Field Identification Guide. 1–168. doi: https://doi.org/10.6084/m9.figshare.7333838.v1.

Steinberg, D. K., Carlson, C. A., Bates, N. R., Johnson, R. J., Michaels, A. F., and Knap, A. H. (2001). Overview of the US JGOFS Bermuda Atlantic Time-series Study (BATS): A decade-scale look at ocean biology and biogeochemistry. Deep Sea Res 2 Top Stud Oceanogr 48, 1405–1447. doi: 10.1016/S0967-0645(00)00148-X.

Steinberg, D. K., Lomas, M. W., and Cope, J. S. (2012). Long-term increase in mesozooplankton biomass in the Sargasso Sea: Linkage to climate and implications for food web dynamics and biogeochemical cycling. Global Biogeochem Cycles 26. doi: 10.1029/2010GB004026.

Stuhldreier, I., Sánchez-Noguera, C., Roth, F., Cortés, J., Rixen, T., and Wild, C. (2015). Upwelling increases net primary production of corals and reef-wide gross primary production along the pacific coast of costa rica. Front Mar Sci 2. doi: 10.3389/fmars.2015.00113.

Sutherland, M. G., McLean, S. J., Love, M. R., Carignan, K. S., and Eakins, B. W. (2014). Digital Elevation Models of Bermuda : Data Sources, Processing and Analysis. Boulder.

Suzuki, A., and Kawahata, H. (2003). Carbon budget of coral reef systems: An overview of observations in fringing reefs, barrier reefs and atolls in the Indo-Pacific regions. Tellus B Chem Phys Meteorol 55, 428–444. doi: 10.1034/j.1600-0889.2003.01442.x.

Tanzil, J. T. I., Brown, B. E., Dunne, R. P., Lee, J. N., Kaandorp, J. A., and Todd, P. A. (2013). Regional decline in growth rates of massive Porites corals in Southeast Asia. Glob Chang Biol 19, 3011–3023. doi: 10.1111/gcb.12279.

Towle, E. K., Enochs, I. C., and Langdon, C. (2015). Threatened Caribbean coral is able to mitigate the adverse effects of ocean acidification on calcification by increasing feeding rate. PLoS One 10, 139398. doi: 10.1371/journal.pone.0123394.

Tribollet, A., Godinot, C., Atkinson, M., and Langdon, C. (2009). Effects of elevated pCO2 on dissolution of coral carbonates by microbial euendoliths. Global Biogeochem Cycles 23, 2006–2009. doi: 10.1029/2008GB003286.

Turner, J. A., Babcock, R. C., Hovey, R., and Kendrick, G. A. (2017). Deep thinking: A systematic review of mesophotic coral ecosystems. ICES Journal of Marine Science 74, 2309–2320. doi: 10.1093/icesjms/fsx085.

Wilkinson, C. (2008). Status of coral reefs of the world: 2008. Townsville.

Wilkinson, C. R. (1999). Global and local threats to coral reef functioning and existence: Review and predictions. Mar Freshw Res 50, 867–878. doi: 10.1071/MF99121.

Williams, G. J., Sandin, S. A., Zgliczynski, B. J., Fox, M. D., Gove, J. M., Rogers, J. S., et al. (2018). Biophysical drivers of coral trophic depth zonation. Mar Biol 165. doi: 10.1007/s00227-018-3314-2.

Wilson, R. W., Millero, F. J., Taylor, J. R., Walsh, P. J., Christensen, V., Jennings, S., et al. (2009). Contribution of fish to the marine inorganic carbon cycle. Science (1979) 323, 359–362. doi: DOI: 10.1126/science.1157972.

Wolanski, E., Colin, P., Naithani, J., Deleersnijder, E., and Golbuu, Y. (2004). Large amplitude, leaky, island-generated, internal waves around Palau, Micronesia. Estuar Coast Shelf Sci 60, 705–716. doi: 10.1016/j.ecss.2004.03.009.

Yeakel, K. L., Andersson, A. J., Bates, N. R., Noyes, T. J., Collins, A., and Garley, R. (2015). Shifts in coral reef biogeochemistry and resulting acidification linked to offshore productivity. Proc Natl Acad Sci U S A 112, 14512–14517. doi: 10.1073/pnas.1507021112.

